# Fine Tuning Rigid Body Docking Results Using the Dreiding Force Field: A Computational Study of 36 Known Nanobody-Protein Complexes

**DOI:** 10.1101/2023.04.18.537388

**Authors:** Aysima Hacisuleyman, Burak Erman

## Abstract

This paper aims to understand the binding strategies of a nanobody-protein pair by studying known complexes. Rigid body protein-ligand docking programs produce several complexes, called decoys, which are good candidates with high scores of shape complementarity, electrostatic interactions, desolvation, buried surface area, and Lennard-Jones potentials. It is not known which decoy represents the true structure. We studied thirty-seven nanobody-protein complexes from the Single Domain Antibody Database, sd-Ab DB, http://www.sdab-db.ca/. For each structure, a large number of decoys are generated using the Fast Fourier Transform algorithm of the software ZDOCK. The decoys were ranked according to their target protein-nanobody interaction energies, calculated by using the Dreiding Force Field, with rank 1 having the lowest interaction energy. Out of thirty-six PDB structures, twenty-five true structures were predicted as rank 1. Eleven of the remaining structures required Ångstrom size rigid body translations of the nanobody relative to the protein to match the given PDB structure. After the translation the Dreiding interaction (DI) energies of all complexes decreased and became rank 1. In one case, rigid body rotations as well as translations of the nanobody were required for matching the crystal structure. We used a Monte Carlo algorithm that randomly translates and rotates the nanobody of a decoy and calculates the DI energy. Results show that rigid body translations and the DI energy are sufficient for determining the correct binding location and pose of ZDOCK created decoys. A survey of the sd-Ab DB showed that each nanobody makes at least one salt bridge with its partner protein, indicating that salt bridge formation is an essential strategy in nanobody-protein recognition. Based on the analysis of the thirty-six crystal structures and evidence from existing literature, we propose a set of principles that could be used in the design of nanobodies.

## Introduction

Nanobodies form a class of proteins produced by the immune system of camelids and sharks and are the antigen-binding domains of heavy chain-only antibodies also denoted as VHH antibodies. These are small proteins, 12-15 kDa, that have three loops which are the three complementarity determining regions, CDR’s, that play a significant role in the binding of the nanobody with its target proteins^1^. Although nanobodies have evolved to eliminate harmful molecules in the systems of their parenting species, they can be used in applications of protein crystallization, targeting ligands, biosensing and diagnostics.^2,3^ Progress in the use of nanobodies in areas concerning human health is impressive^4-10^. Nanobody technology is rapidly becoming a cornerstone of immunoinformatic as evidenced by the recent integrated database INDI^5^.Computationally, the three basic challenges of nanobody design are to (i) choose the best binding nanobody from a given library that will bind to a given protein, (ii) perform mutations in the CDR regions of a given nanobody to change its binding capacity to a protein, (iii) find the binding location and strength of a given nanobody to a given protein. Computationally, these challenges are most commonly investigated by algorithms majority of which are based on the assumption of rigid body docking. The advances in rigid body docking algorithms are remarkable^11-23^. For a given protein and nanobody, the Fast Fourier Transform algorithm (FFT) generates large numbers of their bound forms, called decoys, which are good binders obtained by scoring functions based on shape complementarity, electrostatic interactions, desolvation, buried surface area, Lennard-Jones potentials, etc. The search algorithm common to all these studies searches exhaustively the entire rotational and translational space of the ligand with the receptor being fixed. In rotational search, the three Euler angles for the ligand are changed by small increments, ca 5-15°, and for every rotation, the translational space defined by a set of grids on the surface of the protein, with grid spacing ca 0.5-1.2Å, is explored using FFT. As a result of this technique, a large number of candidate complexes, decoys, are generated, where the nanobody is bound to different suitable locations on the protein and ranked according to the interaction energy criterion mentioned. The main question to be answered is ‘which one of these decoys is the true solution?’, i.e., which one matches the crystal structure? In the present study, by examining 36 known nanobody-protein crystal structures, we see that the dominant metric that serves as best distinguishing feature is the interaction energy calculated by the Dreiding force field^24^ which emerges as an excellent criterion for identifying the decoy that is closest to the known crystal structure. We generate the decoys by the algorithm of ZDOCK^11^. These structures are already good candidates that passed the ZDOCK criteria of docking. We then calculate their interaction energies using the Dreiding potential and rank these candidates with respect to their Dreiding interaction (DI) energies, the decoy with the lowest interaction energy being of rank 1. Then, we find the rank of the decoy whose structure is closest to that of the known structure. The known crystal structures are taken from the data bank Single Domain Antibody Database^25^ (sd-Ab DB), http://www.sdab-db.ca/. As will be shown in detail below, the prediction of the correct nanobody-protein complex using the DI energy as the force field is extremely accurate and reliable. In several cases, the method predicts the lowest DI energy decoy, i.e., rank 1 structure, as the known crystal structure. In some other cases, the decoy that matches the crystal structure is not of rank 1, but small, sub-Ångstrom, rigid body displacements superpose them on the crystal structure. The decoy becomes rank 1 after these rigid body translations. Only for one case, both rigid body translation and rotation are required for the lowest root mean square deviation(RMSD) yielding decoy to become of rank 1. For this purpose, we developed a Monte Carlo script for randomly translating and rotating a decoy until its DI energy reaches a minimum.

The aim of this paper is to learn the binding strategies of nanobody-protein complex formation, with the hope of using them for predicting unknown nanobody-protein systems in the future. There are already several studies that aim at uncovering the rules of recognition. Notably, Gray et al^26^ introduced the concept of binding funnel formation which is essentially an energy funnel into which the ligand is trapped prior to finding its exact location on the protein. The concept has its roots in Hill’s theory ^27^ of cooperativity in biochemistry where the ligand anchors itself on the surface of the protein and exhibits fluctuations in the three Euler angles formed between the ligand and the protein. While the Hill model was for rigid bodies, Gray et al^26^ added side-chain flexibility to small translations and rotations of the ligand as was also done by Fernandez-Recio et al. ^28^, Parma et al. ^29^. Flexibility was also introduced by relaxing interaction potentials by Chen and Weng^30^. Our study shows that once the proper decoy is obtained, small rigid body translations and rotations that lower the interaction energy are sufficient to make it the top-ranking decoy. The implication of this observation is that perfect jigsaw like matching is the dominant potential for defining the best binding pose. The concept of jigsaw matching has indeed been employed in earlier studies of recognition both at the intramolecular and intermolecular level^31-34^. A survey of the sd-Ab DB showed that, with the exception of one or two cases, each nanobody makes at least one salt bridge with its partner protein. Thus, the binding strategy that emerges from our study has two simple ingredients: First, the nanobody anchors itself on the protein at a location that leads to a salt bridge. Then the ligand performs small rigid body translations and rotations about this anchor point until the binding energy is minimized. The rapid formation of a salt bridge between the ligand and the protein as a determinant of recognition has already been acknowledged^35, 36^. The role of flexibility as a nanobody-protein recognition strategy has yet to be seen.

## Methods

We used the Single Domain Antibody Database^25^, http://www.sdab-db.ca/, for obtaining the nanobody-protein crystal structures. Currently, there are 195 Protein Data Bank(PDB) entries in sd-AB database. However, some of these are for single nanobody structures, some are for the same protein with small differences of the nanobodies, and some have missing parts. We eliminated those and chose 36 complexes as our data set.

In the first stage of calculations, we used the FFT algorithm of the software ZDOCK^11^, version 2.3.2 to generate the decoys. Each decoy had a different docking pose. Generally, a set of 100 decoys were sufficient to include the structure closest to the known crystal structure. In only two cases, 4CDG and 4AQ1, more than 100 decoys were necessary as we reported below. Among the various docking programs, we specifically chose ZDOCK because we found it extremely fast, accurate and user-friendly. The method of generating these poses is described in detail in the original paper by Chen and Weng^30^. The algorithm rotates the ligand around the centrally fixed protein by 15 degrees, around each coordinate axis. Following each rotation, it translates the ligand, using the FFT algorithm, along grids with resolution of 1.2 Å constructed on the surface of the protein. We then calculated the DI energy^24^ for every decoy, using the software Biovia Discovery Studio program^37^, which is obtained as the difference between the energy of the complex and the individual energies of unbound protein and nanobody.

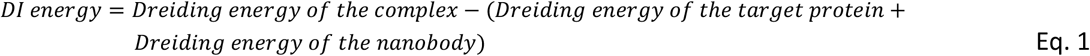

All reported DI energies are in kJ/mol. The protein and the nanobody are taken as in their crystal PDB structures in the absence of water and hydrogens are removed. The DI energy for each decoy calculated in this manner made it possible for us to rank the output structures. The lowest energy decoy is rank 1, the second lowest is rank 2, etc. We then superposed each decoy on the known crystal structure and calculated the RMSD between the two and chose the lowest RMSD as the *best matching complex*. RMSD’s are calculated using VMD’s *measure RMSD* function.^38^ The ranks and RMSD’s of the best matching complexes are reported in Table 1. We also developed a script that translates and rotates a nanobody around a proposed binding pose and calculates the Dreiding energy. The input data for this script are the PDB structures of the nanobody and the protein, the magnitude of the translation of the nanobody, the location of the nanobody atom around which the nanobody will rotate and the amplitudes of translation and the three Euler angles. For the Monte Carlo scheme, the amplitudes are then multiplied by a uniform random number, *r*, in the interval, −1 ≤ *r* ≤ 1, and the nanobody is translated and rotated by the random amount at each step. As will be discussed in detail in the discussion section, once the best fitting complex is found, translating and rotating the nanobody rapidly leads to the conformation of the known crystal structure with the lowest energy. Thus, the translation-rotation process may be applied to several candidate decoys to obtain their true lowest energies.

**Table 1:**
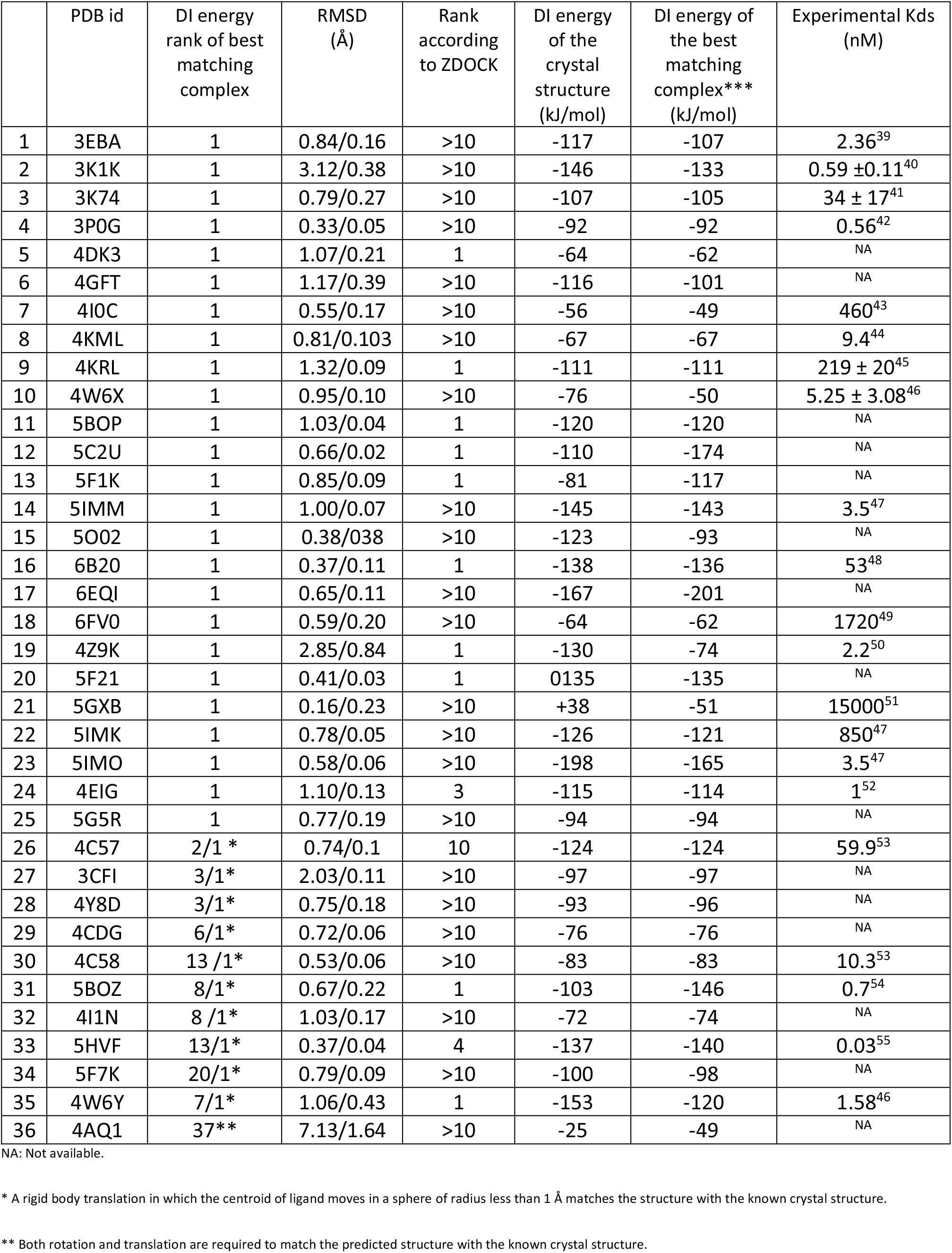
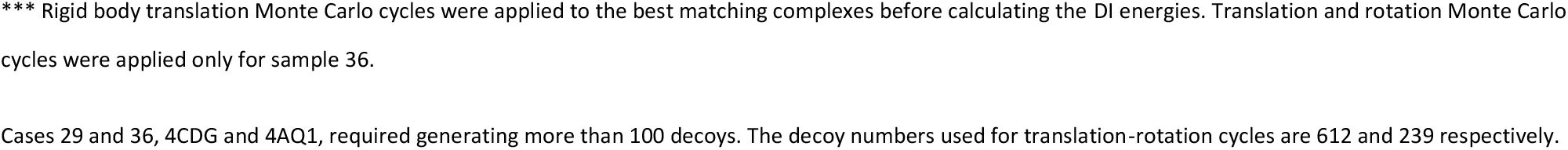
DI energy, RMSD and experimental K_d_ values of 36 selected protein-nanobody complexes.

|  | PDB id | DI energy rank of best matching complex | RMSD (Å) | Rank according to ZDOCK | DI energy of the crystal structure (kJ/mol) | DI energy of the best matching complex*** (kJ/mol) | Experimental K<sub>d</sub>s (nM) |
| --- | --- | --- | --- | --- | --- | --- | --- |
| 1 | 3EBA | 1 | 0.84/0.16 | >10 | -117 | -107 | 2.36<sup>39</sup> |
| 2 | 3K1K | 1 | 3.12/0.38 | >10 | -146 | -133 | 0.59 ± 0.11<sup>40</sup> |
| 3 | 3K74 | 1 | 0.79/0.27 | >10 | -107 | -105 | 34 ± 17<sup>41</sup> |
| 4 | 3P0G | 1 | 0.33/0.05 | >10 | -92 | -92 | 0.56<sup>42</sup> |
| 5 | 4DK3 | 1 | 1.07/0.21 | 1 | -64 | -62 | NA |
| 6 | 4GFT | 1 | 1.17/0.39 | >10 | -116 | -101 | NA |
| 7 | 4I0C | 1 | 0.55/0.17 | >10 | -56 | -49 | 460<sup>43</sup> |
| 8 | 4KML | 1 | 0.81/0.103 | >10 | -67 | -67 | 9.4<sup>44</sup> |
| 9 | 4KRL | 1 | 1.32/0.09 | 1 | -111 | -111 | 219 ± 20<sup>45</sup> |
| 10 | 4W6X | 1 | 0.95/0.10 | >10 | -76 | -50 | 5.25 ± 3.08<sup>46</sup> |
| 11 | 5BOP | 1 | 1.03/0.04 | 1 | -120 | -120 | NA |
| 12 | 5C2U | 1 | 0.66/0.02 | 1 | -110 | -174 | NA |
| 13 | 5F1K | 1 | 0.85/0.09 | 1 | -81 | -117 | NA |
| 14 | 5IMM | 1 | 1.00/0.07 | >10 | -145 | -143 | 3.5<sup>47</sup> |
| 15 | 5O02 | 1 | 0.38/0.38 | >10 | -123 | -93 | NA |
| 16 | 6B20 | 1 | 0.37/0.11 | 1 | -138 | -136 | 53<sup>48</sup> |
| 17 | 6EQI | 1 | 0.65/0.11 | >10 | -167 | -201 | NA |
| 18 | 6FV0 | 1 | 0.59/0.20 | >10 | -64 | -62 | 1720<sup>49</sup> |
| 19 | 4Z9K | 1 | 2.85/0.84 | 1 | -130 | -74 | 2.2<sup>50</sup> |
| 20 | 5F21 | 1 | 0.41/0.03 | 1 | 0135 | -135 | NA |
| 21 | 5GXB | 1 | 0.16/0.23 | >10 | +38 | -51 | 15000<sup>51</sup> |
| 22 | 5IMK | 1 | 0.78/0.05 | >10 | -126 | -121 | 850<sup>47</sup> |
| 23 | 5IMO | 1 | 0.58/0.06 | >10 | -198 | -165 | 3.5<sup>47</sup> |
| 24 | 4EIG | 1 | 1.10/0.13 | 3 | -115 | -114 | 1<sup>52</sup> |
| 25 | 5G5R | 1 | 0.77/0.19 | >10 | -94 | -94 | NA |
| 26 | 4C57 | 2/1 * | 0.74/0.1 | 10 | -124 | -124 | 59.9<sup>53</sup> |
| 27 | 3CFI | 3/1* | 2.03/0.11 | >10 | -97 | -97 | NA |
| 28 | 4Y8D | 3/1* | 0.75/0.18 | >10 | -93 | -96 | NA |
| 29 | 4CDG | 6/1* | 0.72/0.06 | >10 | -76 | -76 | NA |
| 30 | 4C58 | 13 /1* | 0.53/0.06 | >10 | -83 | -83 | 10.3<sup>53</sup> |
| 31 | 5BOZ | 8/1* | 0.67/0.22 | 1 | -103 | -146 | 0.7<sup>54</sup> |
| 32 | 4I1N | 8 /1* | 1.03/0.17 | >10 | -72 | -74 | NA |
| 33 | 5HVF | 13/1* | 0.37/0.04 | 4 | -137 | -140 | 0.03<sup>55</sup> |
| 34 | 5F7K | 20/1* | 0.79/0.09 | >10 | -100 | -98 | NA |
| 35 | 4W6Y | 7/1* | 1.06/0.43 | 1 | -153 | -120 | 1.58<sup>46</sup> |
| 36 | 4AQ1 | 37** | 7.13/1.64 | >10 | -25 | -49 | NA |

NA: Not available.
\* A rigid body translation in which the centroid of ligand moves in a sphere of radius less than 1 Å matches the structure with the known crystal structure.
\*\* Both rotation and translation are required to match the predicted structure with the known crystal structure.
\*\*\* Rigid body translation Monte Carlo cycles were applied to the best matching complexes before calculating the DI energies. Translation and rotation Monte Carlo cycles were applied only for sample 36.
Cases 29 and 36, 4CDG and 4AQ1, required generating more than 100 decoys. The decoy numbers used for translation-rotation cycles are 612 and 239 respectively.

In addition to energy, RMSD calculations and ranking, we used this dataset to identify the salt bridges in the crystal structures using the electrostatic non-bonded interactions module of Discovery Studio.Identifying the salt bridge which will form between the nanobody, and the protein is crucial for simplifying the problem since it reduces the 12-dimensional phase space available to the complex to three dimensions defined by the three Euler angles.

## Results

### (i) Energy rank of the nearest-native structure

The list of structures, the energy rank of the best matching complex and RMSD between the decoy and the corresponding crystal structure are given in columns 2-4 of Table 1. A rank of 1 in column 3 indicates that the best matching complex to the crystal structure, i e., the lowest RMSD, has the lowest DI energy among all the decoys generated. Column 3 in Table 1 shows that 25 structures have rank 1, indicating that the decoy closest to the crystal structure already has the lowest DI energy. The closeness to the crystal structure is given by the RMSD values in column 4. Entries from 26 to 35 in Table 1 show predictions which do not have rank 1, i.e., the DI energy is not the lowest for these predictions. However, the three-dimensional structures of these structures are close to those of the corresponding crystal structures and may be made to coincide by pure rigid body translation less than an Ångstrom (see below). In the fifth column of Table 1, we compared the ranks obtained using the scoring function 3.0.2 of the ZDOCK server, which is based on surface complementarity, electrostatic interactions, desolvation, buried surface area, and various intermolecular potentials as mentioned in the introduction. The ZDOCK 3.0.2 returns the decoys in a sorted order from highest score being the decoy having the lowest energy to the lowest score decoys having higher energies. A value of 10> in column 5 indicates that the best matching complex is not among the 10 highest score decoys given by the ZDOCK 3.0.2 scoring system. Column 5 shows that 24 of the 36 complexes are not in the set of first 10 decoys according to the ZDOCK 3.0.2 scoring system. Entry 36 in Table 1 required both translation and rotation for a perfect fit with crystal structure.

The Dreiding energy obtained from the crystal structure is given in column 6. Column 7 shows the Dreiding energy for the best matching complex after rigid body translation Monte Carlo cycles. Column 8 shows the available *K*_*D*_ values. The superscript following each entry in column 8 indicates the reference from which the data is taken. Columns 6 and 7 may be used in determining good binders of unknown complexes as will be discussed in more detail in the discussion section.

We now give a few representative results from the analysis of the set of 36 known cases:

#### 1. The case of 3EBA

As an example of energy ranking, we present the energy rank plot of 100 decoys of 3EBA in Figure 1. Ordinate values represent the DI energies obtained, data points are for 100 decoys where the filled circle shows the energy of the complex that best fits the crystal structure of 3EBA. The empty circles represent results for the remaining decoys.

**Figure 1:**
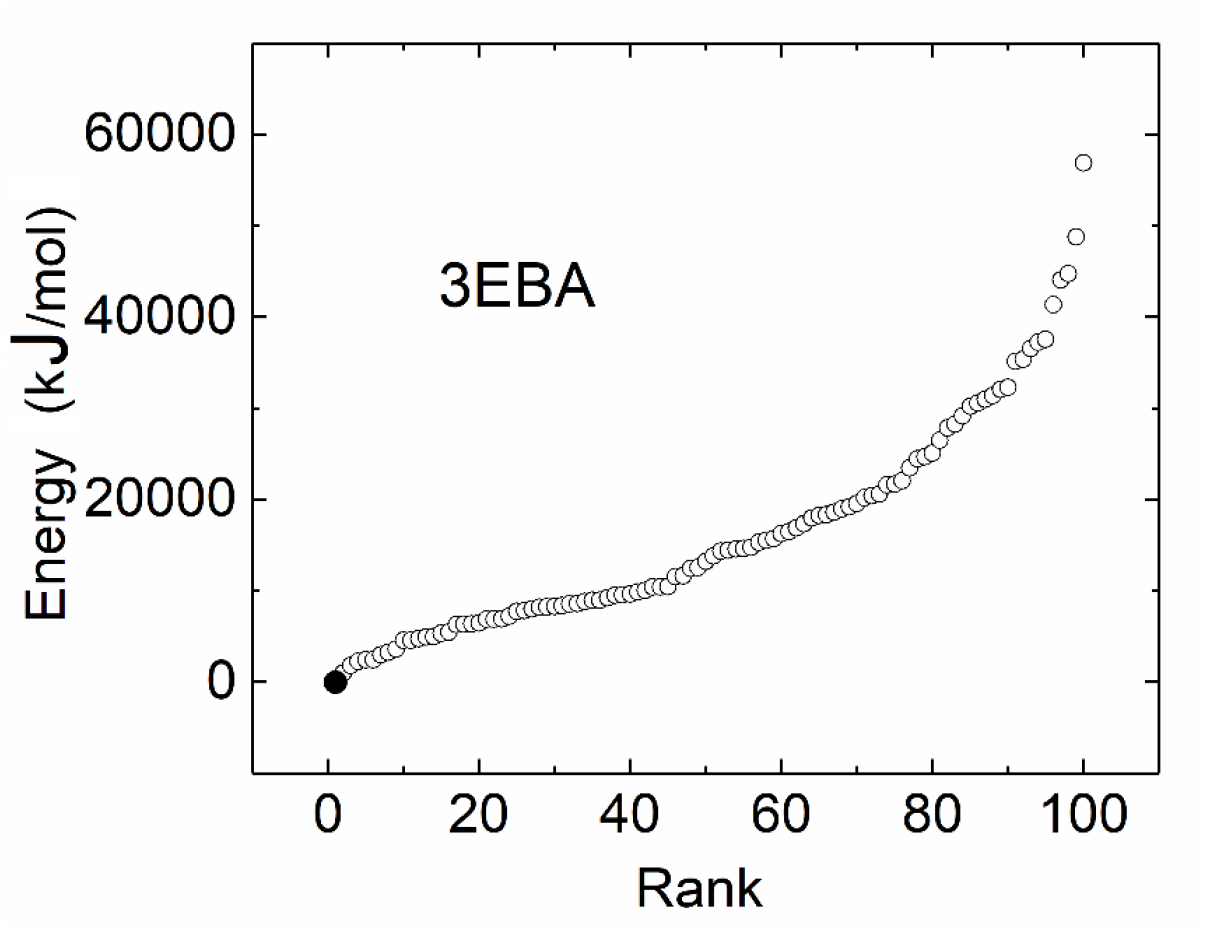
DI energy of 100 decoys from ZDOCK for the protein-nanobody complex, 3EBA, is plotted as a function of the rank, where the filled circle shows the DI energy of the best matching complex, meaning the decoy having the lowest RMSD from the crystal structure.

We see that the best matching complex has the lowest DI energy, meaning its rank is 1. The intermolecular hydrogen bond strength calculated by using Discovery Studio for the decoy with rank 1 is the strongest among all other decoys. However, this is not a general trait, and we cannot conclude that all rank 1 structures have the strongest hydrogen bonding.

#### 2. The case of 3CFI

As another example, we present the energy rank plot of 3CFI decoys in Figure 2. Column 3 for 3CFI entry in Table 1, 3/1*, indicates that the prediction closest to crystal structure had energy rank 3 but small rigid body translations resulted in almost perfect superposition with crystal structure and improved the DI energy to rank 1. The entry 2.03/0.46 in column 4 of Table 1 for 3CFI indicates that the RMSD of rank 3 decoy, the filled circle, was 2.03 Å and dropped to 0.46 Å after a rigid body translation and the value of its DI energy decreased to -99 kJ/mol upon the rigid body translation. Rigid body translation was applied to several decoys of low energy of Figure 2, but none went below -99 kJ/mol. The next lowest binding energy was -42 kJ/mol. Thus, rigid body translation decreases the energies, but none goes below that of the best matching structure. It should be added that this energy, although negative, does not represent the true energy because the contribution of water and hydrogens are missing.

**Figure 2:**
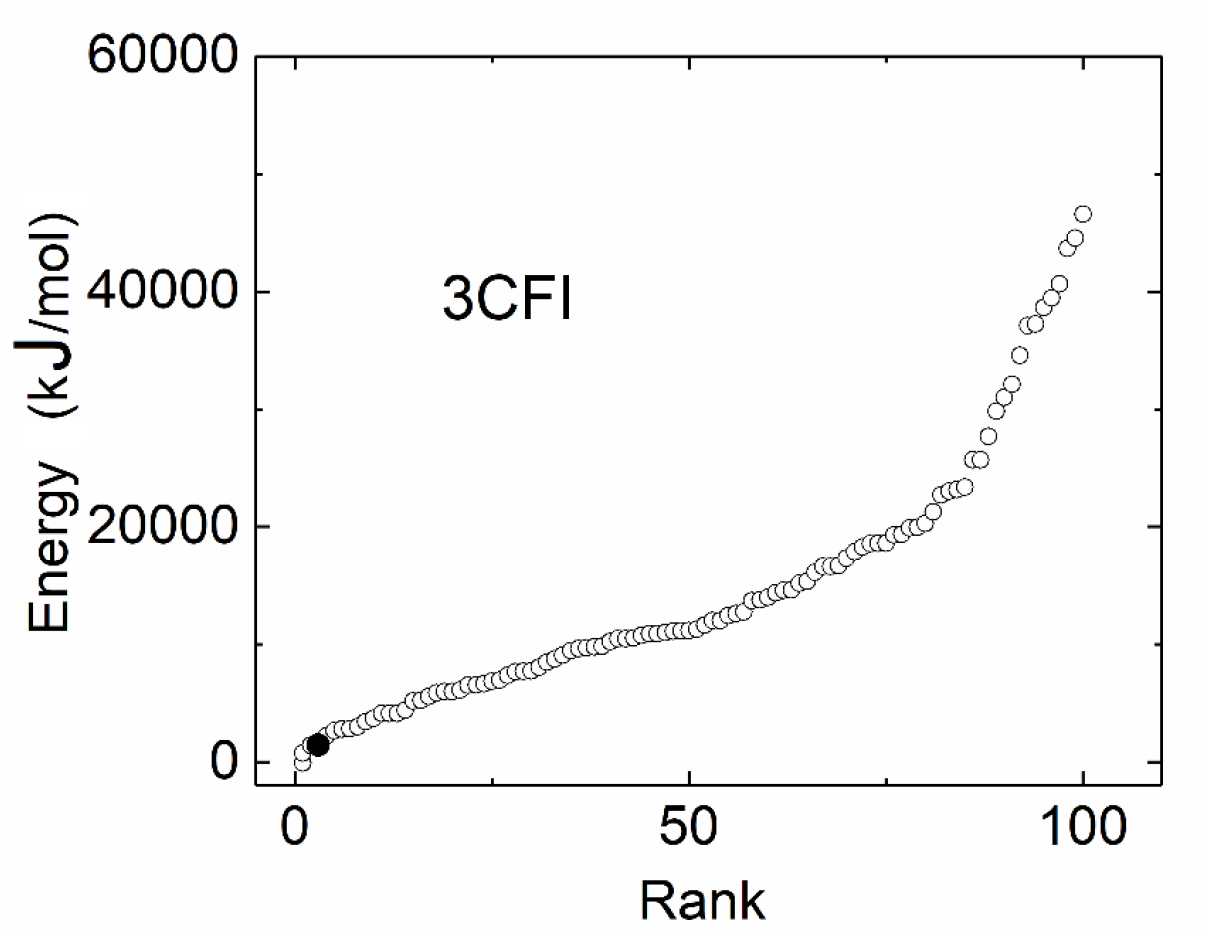
DI energy of 100 decoys from ZDOCK for the protein-nanobody complex, 3CFI, is plotted as a function of the rank, where the filled circle shows the DI energy of the best matching complex, meaning the decoy having the lowest RMSD from the crystal structure.

The 10 decoys indicated by a single asterisk in column 3 of Table 1, have relatively low RMSD’s as shown by the values to the left of the slashes in column 4 and can be aligned by small rigid body translation of the centroid within a sphere of radius 1 Å. As a result of this operation, the DI energy then becomes rank 1 for each, with RMSD’s shown on the right side of the slashes. The need to translate is because of the size of the grid resolution in generating the decoys, which is 1.2 Å in the software we used. The translation operation that is needed to overcome the grid resolution effect is simple and straightforward and may be programmed such that the DI energy is recorded for each small step translation and the structure showing the lowest DI energy is accepted. This is the procedure we followed for the entries between 26 and 35 in Table 1. The conformation of the decoy of rank 3 is presented in Figure 3 where the nanobody of the decoy is highlighted in yellow. Translating the nanobody by 1.42 Å as a rigid body result in overlap with crystal structure. A more detailed explanation of the translation operation is explained in the discussion section.

**Figure 3:**
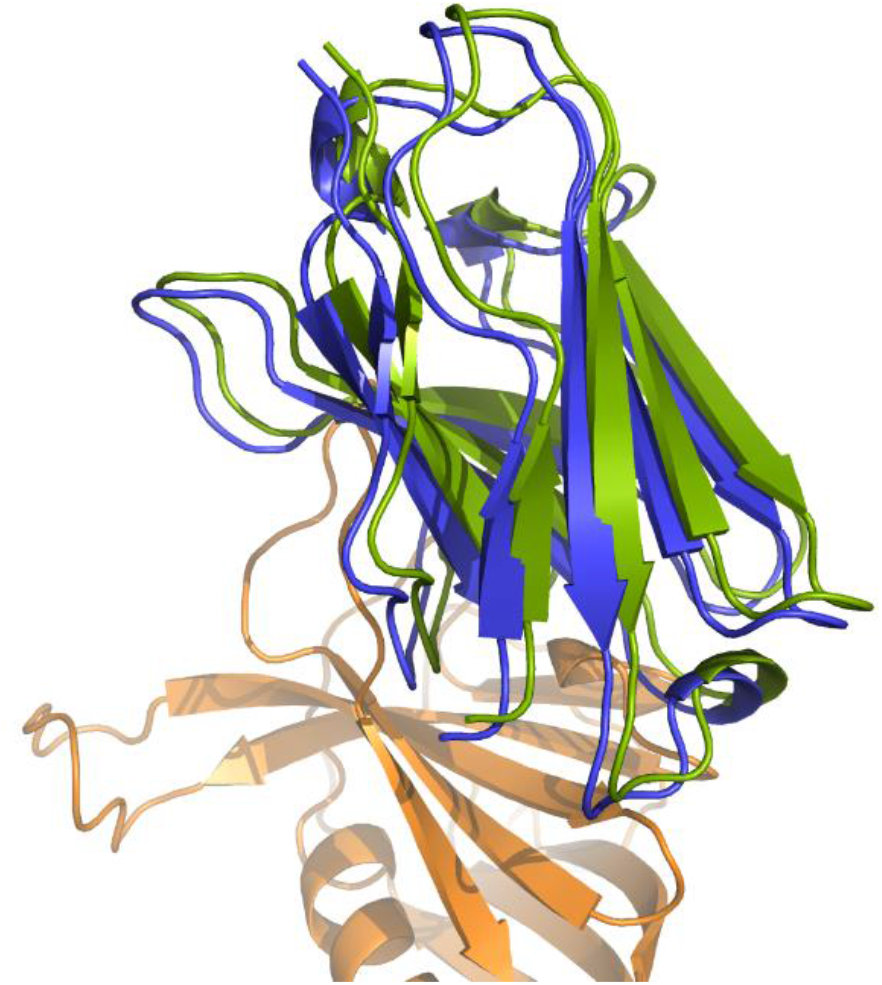
Rank 3 decoy for 3CFI superposed on its crystal structure. Nanobody of the decoy is in green. Crystal structure nanobody is in blue and the target protein is in orange.

#### 3. The case of 4AQ1

The minimum RMSD decoy for 4AQ1 is shown in the left panel of Figure 4 where the nanobody is colored in blue. Due to the size and complexity of the protein, 100 decoys were not sufficient to make the prediction and a set of 300 decoys were used. The decoy that is closest to the crystal structure is the decoy 239 out of 300, with a RMSD of 7.13 Å. When the 300 decoys are ordered with respect to their Dreiding energies, this decoy is of rank 37 and cannot be superposed on the crystal structure by pure translation only, and rotation is also required. There is a salt bridge between GLU560 and ARG45 in decoy 239 where the distance between alpha carbons is 5.38 Å. In the crystal structure, there is a salt bridge between LYS565 of the protein and GLU44 of the nanobody where the distance between the alpha carbons is 9.73 Å. The two neighboring pairs LYS565-GLU44 and GLU560-ARG45 may be seen as nuclei of binding according to which a salt bridge between the nanobody and the protein is established first and the nanobody rotates and translates about the salt bridge until its energy is lowered. We first used Monte Carlo translation-rotation cycles on decoy 239 around GLU560 and ARG45. The structure shown on the right panel in Figure 4 is the structure obtained by rigid body translation and rotation about the nitrogen of ARG45 until its DI energy does not decrease further. This operation requires about 1000 Monte Carlo steps of translation and 1000 steps of rotation. The DI energy at the end of these steps is -49 kJ/mol.

**Figure 4:**
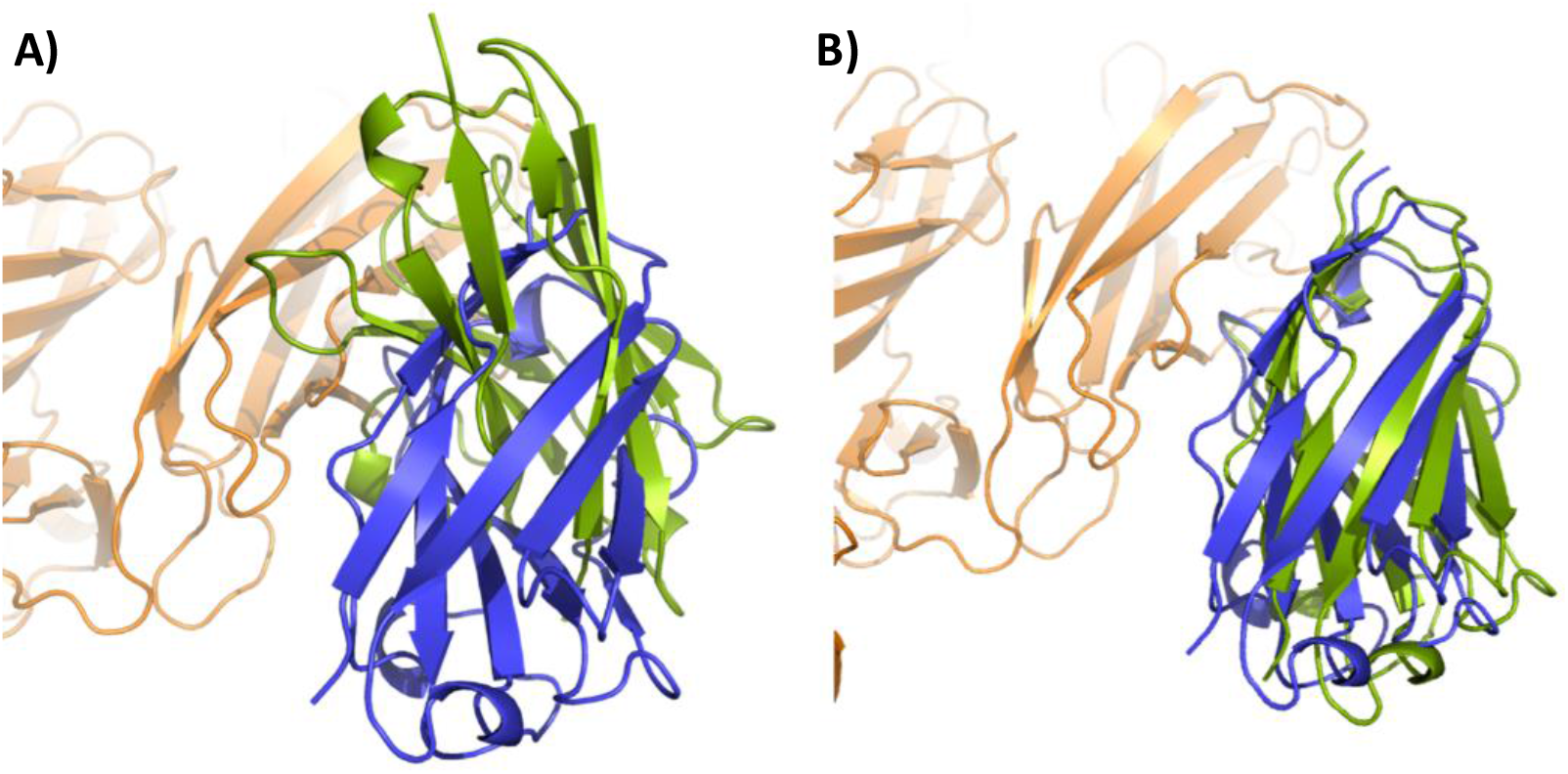
A) Decoy 239 for 4AQI is superposed on 4AQI crystal structure, nanobody of the decoy is highlighted in green and the nanobody from the crystal structure is colored in blue. B) Decoy 239 is superposed on the crystal structure after performing 1000 Monte Carlo steps of translation and 1000 Monte Carlo steps for rotation about the nitrogen atom of ARG45 of the nanobody, where the nanobody is highlighted in green. (See legend to Figure 3 for color scheme)

Secondly, we manually translated the decoy 239 such that GLU44 is in the vicinity of LYS565 of the protein. This translation can be done easily using Discovery Studio such that the side chain Oxygen atom of GLU44 is approximately at a distance of 3 Å from the side chain nitrogen atom of LYS565. We then applied translation-rotation cycles, 2000 Monte Carlo steps each, until a minimum binding energy of -52 kJ/mol was obtained. However, this minimum energy structure deviated from the crystal structure by a RMSD of 1.64 Å shown in Figure 5. The DI energy of the nanobody in the crystal structure is -24 kJ/mol which is higher than the Monte Carlo translated rotated structure. This may not reflect the real picture and (i) the presence of water and/or (ii) deformation of side chains may make the crystal structure more favorable. These two factors have not been thoroughly investigated in the literature. Yet, the conformational deviations due to these two effects are small as verified by Figure 5.

**Figure 5:**
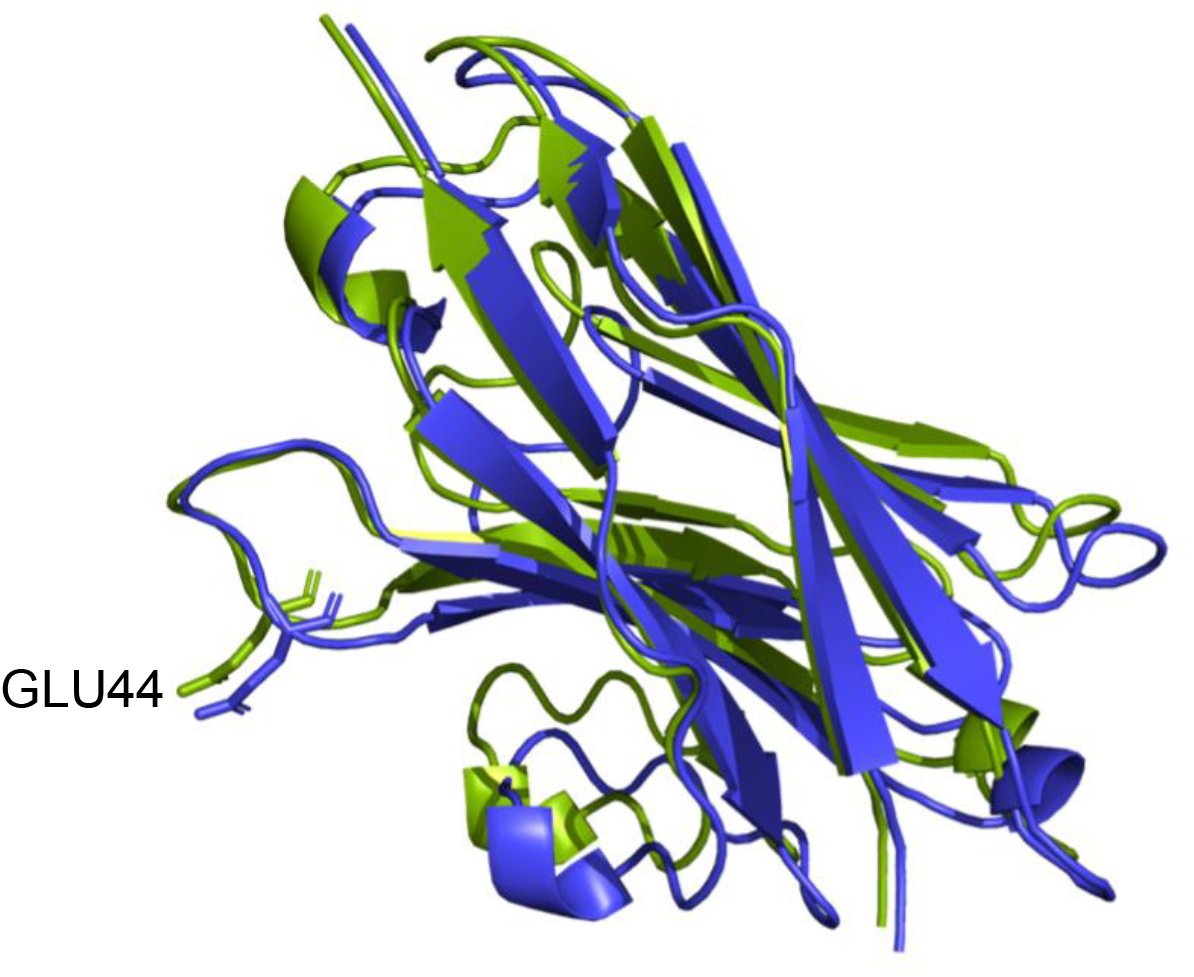
The nanobody of the decoy 239 is superposed on the nanobody crystal structure of 4AQI, after performing a second Monte Carlo translation-rotation cycle of 2000 steps each about the oxygen atom of GLU44 of the nanobody, where the decoy nanobody is colored in green and the nanobody from the crystal structure is colored in blue.

**Figure 6:**
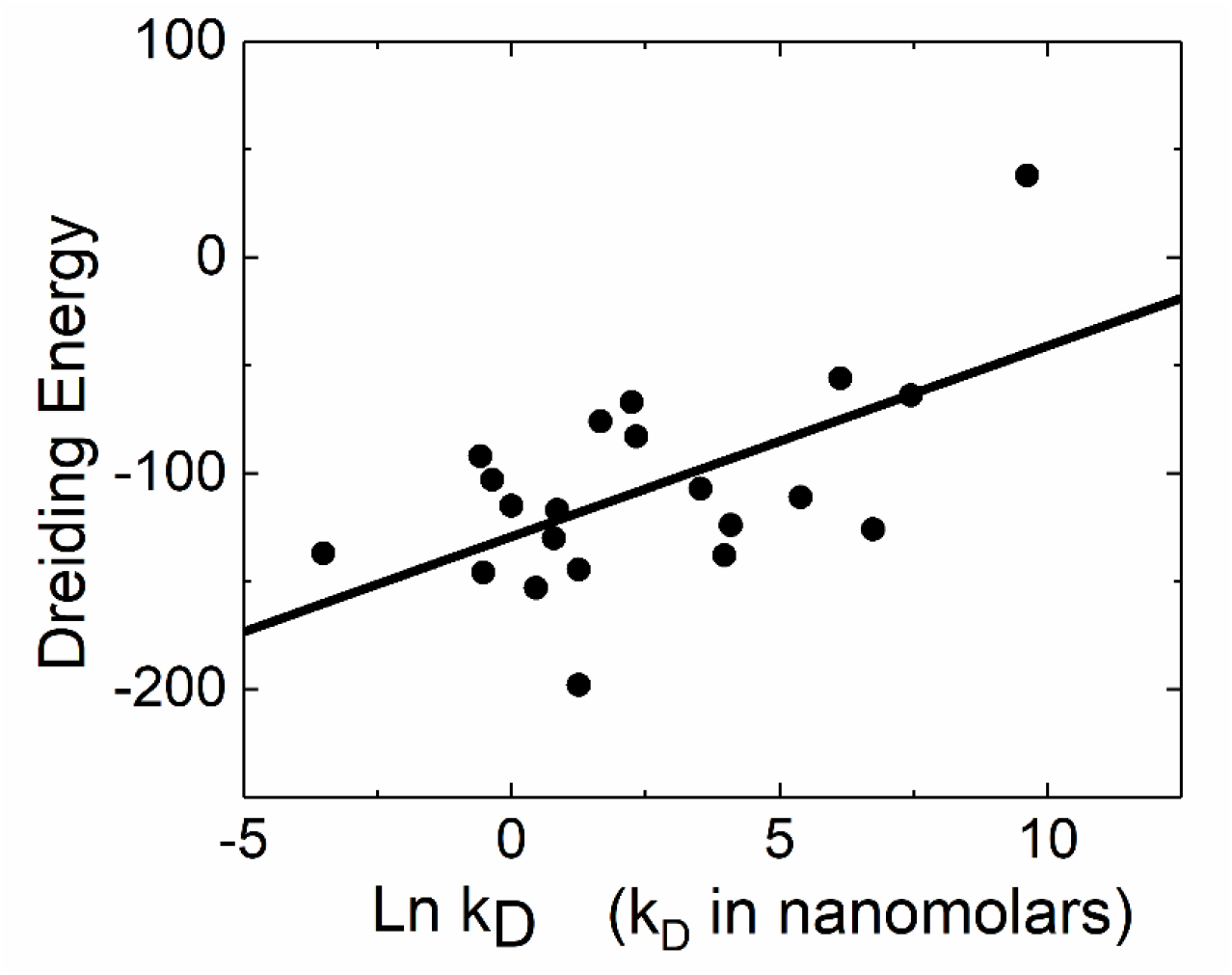
DI energy(kJ/mol) of complexes with available experimental *K*_*D*_ values are plotted against their experimental *lnK*_*D*_(nM) values.

## Discussion

The study of thirty-seven nanobody-protein complexes from the PDB showed that the Dreiding force field is a good metric in the prediction of the crystal structure from among a large number of decoys. For each complex, 100, (300 for one case), decoys were generated by ZDOCK which have favorable shape complementarity, electrostatic interactions, desolvation and buried surface area scores according to the ZDOCK scoring system. Out of thirty-seven PDB structures, twenty-five of them were predicted as rank 1 when ranked according to the Dreiding energy. Eleven of the remaining structures required Ångstrom size rigid body displacements of the nanobody relative to the protein to match the given PDB structure. After the translation the complexes all became rank 1. In one case, neither rigid body rotations nor translations of the nanobody resulted in matching with the crystal structure. This was partly due to the presence of two minima of approximately the same energies. We used a Monte Carlo algorithm that randomly translates and rotates the nanobody of a decoy and calculates the Dreiding energy and the unbiased Monte Carlo simulation did not prefer the crystal structure. This script was applied to the cases that were not rank 1 and sub-Ångstrom size rigid body translation was sufficient for matching.

The observation that all nanobody-protein complexes exhibit at least one salt bridge suggests that salt bridges play essential role on the binding process. Statistical mechanical arguments given earlier^27^ have suggested that binding is initiated by anchoring of the ligand at a suitable site on the protein followed by rotational rearrangements of the ligand around the fixed anchor point. The identification of the salt bridging residue pairs by the ligand constitutes the onset of the binding funnel^26^ in which the nanobody exhibits small translational and rotational rearrangements until it locks into the protein. This idea forms the basis of the Monte Carlo script that we used in the present study. Among the 98 nanobody-protein complexes whose crystal structures are listed in sd-Ab DB and listed in the supplementary Material Table S1, all but five complexes exhibit at least one salt bridge. The reason why five complexes do not have salt bridges is because there are no charged residues in the three CDR loops of the nanobodies. Most of the bonding in these five exceptional cases is through polar residues. The distribution of salt bridge pairs with respect to residue type is presented in Table 2.

**Table 2:** Number of salt bridge forming residue pairs from 98 nanobody-protein complexes listed in sd-Ab DB

|  | ASP | GLU |
| --- | --- | --- |
| ARG | 57 | 71 |
| LYS | 33 | 16 |

The C^α^-C^α^ distances of salt bridge forming residues lie in the range 8-12 Å. More importantly, the distance between the donor and acceptor atoms on the side chains is approximately 3 Å. The ZDOCK decoys all exhibit salt bridges, which point to the anchor point around which the nanobody will fluctuate. Fixing the anchor point of the ligand reduces the phase space significantly. Thus, selecting the salt bridge at the beginning of the docking and using Monte Carlo translations and rotations may be a useful simplifying strategy. The total number, *N*_*SB*_, of possible salt bridges between a protein and a nanobody is

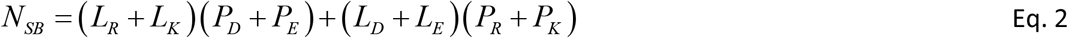

where, *L* and *R* denote the ligand and the protein and the subscripts *R, K, D*, and *E* denote Arg, Lys, Asp and Glu, respectively, on the right hand side of Eq.1. Of course, the ZDOCK decoy predictions followed by DI energy calculations eliminate most of the *N*_*SB*_ structures and the prediction algorithm may reduce to only few Monte Carlo simulations around each of the designated candidate decoy.

The DI energies of the complexes with available experimental *K*_*D*_ values, column 7 of Table 1 are plotted as a function of available *K*_*D*_ ‘s, column 8. The solid line is the linear fit through the points given by the equation

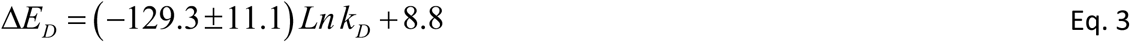

where Δ*E*_*D*_ is the DI binding energy. Solving for *K*_*D*_ leads to

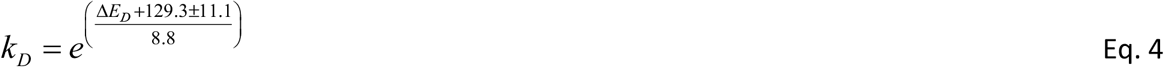

Equation 3 is plotted in Figure 7 where the ordinate is in logarithmic scale and the abscissa values span the range −50 ≤ Δ*E*_*D*_ ≤ −200 kJ/mol which covers the scale from poor to excellent binding. This figure may be used as a master chart to assess the quality of binding in unknown nanobody-protein pairs. The DI energy obtained from the Monte Carlo translation-rotation operation on low-rank decoys will give the corresponding *K*_*D*_ values.

**Figure 7:**
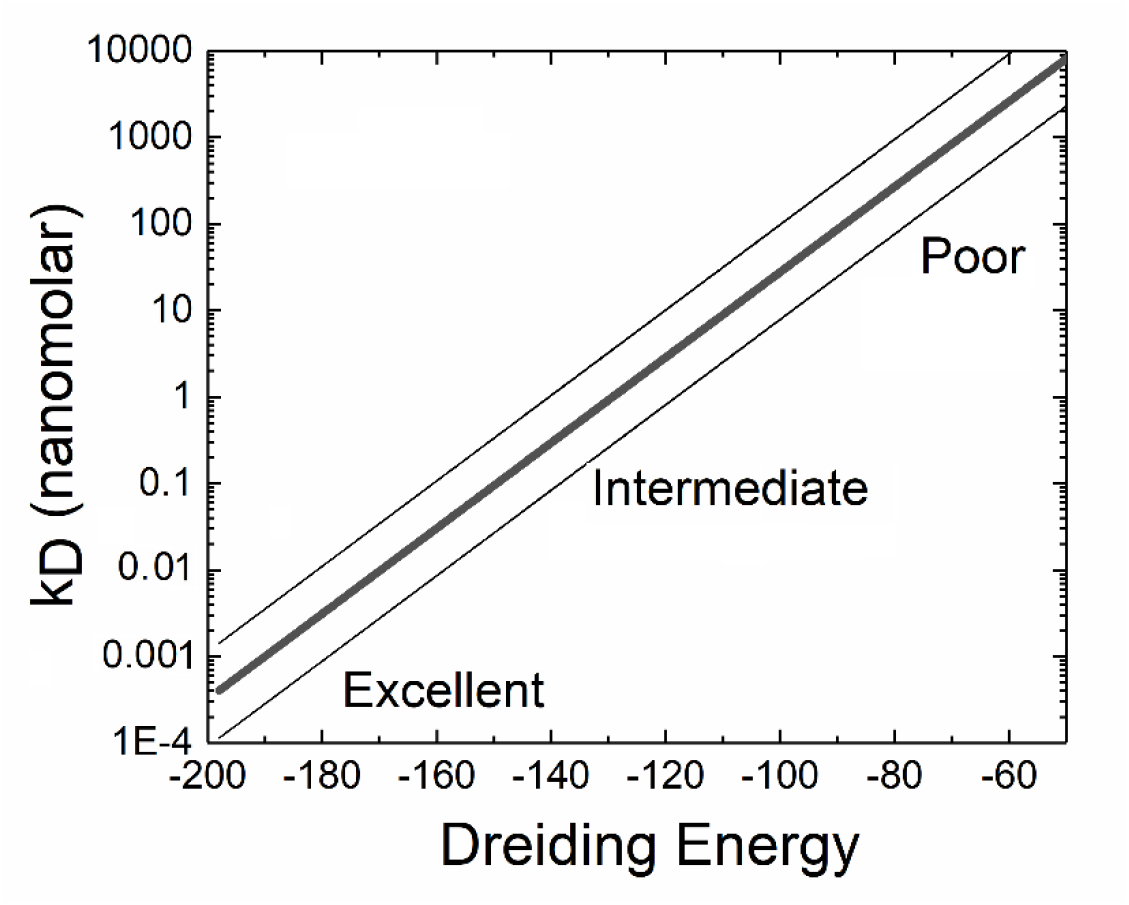
The *K*_*D*_ values are plotted against the DI energy using Eq. 4. Ordinate scale is logarithmic. *K*_*D*_ of a given protein-nanobody complex whose DI energy is known can be obtained from this figure.

The findings of the present work may be used as design principles in nanobody engineering. We now discuss two cases:

*Case 1:* Crystal structures of the nanobody and the protein are known but the structure of the complex is not known. We run a ZDOCK simulation and obtain several decoys, one of which is possibly the true solution. We determine the DI energies of all the decoys and rank them. Starting from the lowest rank, we perform Monte Carlo translation-rotation simulations and obtain the lowest DI energy decoy. This will possibly be the structure closest to the true structure.

*Case 2:* The structure of the complex is known, and modifications are required to increase the binding affinity: We repeat the steps of Case 1 for each mutation and obtain the binding pose and the DI energy of the structure closest to the true structure.

The bottleneck in the above workflow is the Monte Carlo step. In an ordinary laptop, 2000 step Monte Carlo cycle of diverse structures and the determination of the DI energy is in the order of one hour.

