## Supplemental Table 1 for "Fine Tuning Rigid Body Docking Results Using the Dreiding Force Field: A Computational Study of 36 Known Nanobody-Protein Complexes"

Table S1: List of salt-bridge forming nanobody residue pairs from the Single Domain Antibody Database, sd-Ab DB, <http://www.sdab-db.ca/> for nanobody-protein complexes. The nanobody is identified with C and the protein with B. Cases without entry indicate the absence of salt bridge.

| 5o02.pdb | C:ARG99 B:GLU486 |
| --- | --- |
| 5o04.pdb | C:LYS96 B:PHE532 |
| 5o0w.pdb | C:ARG123 B:GLU245 |
| 5ocl.pdb | C:ASP37 B:ARG74 |
| 5omn.pdb | C:ASP54 B:ARG484  C:GLU64 B:ARG492  C:ARG98 B:GLU496 |
| 5ovw.pdb | C:ASP52 B:LYS223  C:ARG123 B:GLU245 |
| 5tjw.pdb | C:ARG53 B:GLU375  C:ASP56 B:LYS103 |
| 5uk4.pdb | C:LYS23 B:ASP359  C:ARG27 B:GLU377  C:ARG54 B:ASP384  C:LYS100 B:ASP374 |
| 5uz7.pdb | C:ARG98 B:GLU226 |
| 5vak.pdb | C:ARG72 B:ASP220 |
| 5vxj.pdb | C:ARG19 B:ASP97  C:ASP73 B:LYS137  C:LYS76 B:GLU122 |
| 5vxk.pdb | C:ARG105 B:GLU288 |
| 5vxl.pdb | C:ASP52 B:LYS205  C:ARG102 B:GLU201 |
| 5vxm.pdb | C:ASP53 B:LYS205  C:ASP54 B:LYS205  C:ARG59 B:GLU201 |
| 6b20.pdb | C:ARG99 B:ASP228 |
| 6b73.pdb | C:ASP62 B:LYS91  C:ASP73 B:ARG252 |
| 6c9w.pdb | C:ARG52 B:PHE250 |
| 6eqi.pdb | C:ARG27 B:ASP303  C:ARG53 B:ASP338  C:ARG97 B:GLU299 |
| 6fv0.pdb | --- |
| 3cfi.pdb | C:ARG53 B:ASP85  C:ASP103 B:LYS110  C:ARG107 B:GLU135 |
| 3k1k.pdb | C:ARG35 B:GLU142  C:GLU44 B:LYS166  C:ARG56 B:GLU172  C:GLU101 B:ARG168 |
| 3g9a.pdb | C:ASP95 B:LYS166  C:ASP101 B:LYS166 |
| 3k74.pdb | C:LYS218 B:GLU17 |
| 3p0g.pdb | C:GLU106 B:ARG131 |
| 4c57.pdb | C:GLU58 B:ARG43  C:ARG107 B:GLU56 |
| 4c58.pdb | C:ARG34 B:GLU308  C:ARG72 B:GLU315 |
| 4dk3.pdb | C:GLU105 B:ARG703 |
| 4eig.pdb | C:LYS33 B:GLU17 |
| 4eiz.pdb | C:LYS112 B:ASP11 |
| 4fhb.pdb | C:ARG101 B:ASP127  C:LYS111 B:GLU118 |
| 4gft.pdb | C:ARG29 B:GLU179  C:ARG101 B:GLN204 |
| 4hep.pdb | --- |
| 4i0c.pdb | C:PHE55 B:ARG122 |
| 4i13.pdb | C:ARG100 B:GLU48  C:ARG104 B:GLU17  C:LYS113 B:ASP69 |
| 4i1n.pdb | C:ARG100 B:GLU48  C:LYS113 B:ASP69 |
| 4kdt.pdb | C:ARG31 B:GLU44  C:ARG50 B:GLU44  C:ASP56 B:ARG81  C:ARG105 B:GLU47  C:ASP106 B:LYS41  C:LYS110 B:GLU77 |
| 4kml.pdb | C:ASP59 B:HIS177  C:ARG106 B:HIS177 |
| 4krl.pdb | C:ARG30 B:ASP355  C:GLU110 B:ARG353  C:ASP112 B:ARG353 |
| 4krp.pdb | C:ARG27 B:GLU431  C:ASP111 B:ARG310  C:GLU113 B:ARG405  C:ASP115 B:ARG403 |
| 4ldo.pdb | C:ASP99 B:LYS1267  C:ASP106 B:ARG1131 |
| 4lgp.pdb | C:ARG52 B:HIS94  C:ARG104 B:GLU102 |
| 4lgr.pdb | C:PHE105 B:ARG125 |
| 4lgs.pdb | C:ASP28 B:ARG125  C:ARG31 B:HIS94 |
| 4lhj.pdb | C:GLU32 B:ARG235 |
| 4lhq.pdb | C:ARG31 B:HIS94 |
| 4mqs.pdb | C:GLU109 B:ARG135 |
| 4n9o.pdb | C:ARG106 B:HIS177  C:ASP59 B:HIS177 |
| 4pir.pdb | C:ARG105 B:GLU173  C:ARG110 B:GLU173 |
| 4qkx.pdb | C:ASP101 B:LYS1267  C:ASP108 B:ARG1131 |
| 4s10.pdb | C:ARG230 B:ASP99 |
| 4w2q.pdb | C:ARG30 B:ASP679 |
| 4w6x.pdb | C:ASP114 B:LYS150 |
| 4w6y.pdb | C:GLU101 B:LYS114  C:ARG108 B:GLU96  C:ARG112 B:GLU122 |
| 4wgv.pdb | C:ARG53 B:ASP290  C:ARG73 B:ASP296  C:ARG100 B:ASP290 |
| 4x7c.pdb | C:LYS96 B:PHE524 |
| 4x7e.pdb | C:LYS96 B:PHE532 |
| 4x7f.pdb | C:GLU54 B:ARG501 |
| 4xt1.pdb | C:GLU104 B:ARG225  C:ASP106 B:ARG221 |
| 4y8d.pdb | C:ARG30 B:GLU308  C:ARG62 B:GLU315  C:GLU104 B:ARG164 |
| 4z9k.pdb | C:ASP53 B:ARG126 |
| 5bop.pdb | C:ARG99 B:ASP152 |
| 5boz.pdb | C:ARG27 B:GLU67  C:ARG103 B:GLU138  C:ASP112 B:ARG166  C:ASP114 B:ARG166 |
| 5c1m.pdb | C:ARG21 B:ASP177  C:ASP62 B:LYS100  C:ASP66 B:LYS100  C:ASP73 B:ARG258 |
| 5c2u.pdb | C:ASP26 B:LYS322  C:ARG50 B:GLU294  C:ARG96 B:GLU315  C:ARG97 B:GLU315 |
| 5da0.pdb | C:ASP35 B:ARG463  C:GLU47 B:ARG483 |
| 5dfz.pdb | --- |
| 5f1k.pdb | C:ARG27 B:GLU292  C:ARG100 B:GLU292  C:ARG106 B:ASP202 |
| 5f1o.pdb | C:ASP106 B:LYS69  C:ARG28 B:ASP141  C:ARG59 B:GLU72  C:ARG103 B:ASP147 |
| 5f21.pdb | C:LYS56 B:GLU292  C:LYS60 B:ILE300  C:ASP109 B:LYS276 |
| 5f7k.pdb | C:ARG98 B:HIS369 |
| 5f7l.pdb | C:HIS45 B:ARG139  C:ASP100 B:ARG295  C:ASP104 B:ARG295  C:GLU109 B:ARG139 |
| 5foj.pdb | C:LYS54 B:VAL504  C:ARG55 B:ASP371 |
| 5g5r.pdb | C:GLU46 B:ARG331  C:ASP61 B:ARG273  C:ARG104 B:GLU272 |
| 5g5x.pdb | C:GLU46 B:ARG331  C:ASP61 B:ARG273  C:ARG104 B:GLU272 |
| 5gxb.pdb | C:ARG101 B:GLU374 |
| 5hvf.pdb | C:ASP107 B:ARG12 |
| 5hvg.pdb | C:ASP108 B:ARG384 |
| 5imk.pdb | C:ARG51 B:ASP44  C:LYS102 B:GLU95  C:GLU110 B:ARG108 |
| 5iml.pdb | C:LYS102 B:GLU95  C:GLU110 B:ARG108  C:ARG51 B:ASP44 |
| 5imm.pdb | C:ARG51 B:ASP43  C:LYS102 B:ASP43  C:GLU110 B:ARG107  C:ASP112 B:LYS109 |
| 5imo.pdb | C:ARG51 B:ASP44  C:ASP101 B:ARG40  C:LYS102 B:ASP44  C:GLU110 B:HIS1  C:ASP112 B:LYS110 |
| 5ip4.pdb | C:ARG55 B:ASP294 |
| 5j1s.pdb | --- |
| 5jds.pdb | C:LYS27 B:GLU60  C:ARG32 B:GLU58  C:ASP99 B:ARG113 |
| 5lhn.pdb | C:LYS27 B:GLU60 C:ARG32 B:GLU58  C:ASP99 B:ARG113 |
| 5m14.pdb | C:ARG45 B:ASP41  C:ARG113 B:GLU45 |
| 5m15.pdb | C:ARG45 B:ASP41  C:LYS103 B:GLU111 |
| 5m2i.pdb | C:ARG54 B:ASP140  C:LYS65 B:ASP45 |
| 5m2m.pdb | C:ARG55 B:GLU23  C:LYS70 B:GLU146  C:ARG111 B:GLU23 |
| 5mp2.pdb | C:GLU46 B:GLN49 |
| 5mwn.pdb | C:LYS96 B:GLU120  C:ASP107 B:ARG121 |
| 5my6.pdb | C:GLU44 B:ARG157  C:ASP54 B:LYS150  C:GLU102 B:LYS170 |
| 5mzv.pdb | C:ARG52 B:GLU108  C:LYS65 B:ASP40  C:ARG106 B:ASP63  C:ASP111 B:LYS217 |
| 5lhr.pdb | C:ASP99 B:LYS192  C:ARG104 B:ASP189 |
| 5o2u.pdb | C:ARG50 B:GLU213 |
| 5toj.pdb | --- |
| 6ey0.pdb | C:ARG100 B:GLU143  C:ASP101 B:ARG104 |

`
